# The disease associated Tau35 fragment has an increased propensity to aggregate compared to full-length tau

**DOI:** 10.1101/2021.07.08.451675

**Authors:** Chen Lyu, Stefano Da Vela, Youssra Al-Hilaly, Karen E. Marshall, Richard Thorogate, Dmitri Svergun, Louise Serpell, Annalisa Pastore, Diane Hanger

## Abstract

Tau35 is a truncated form of tau found in human brain in a subset of tauopathies. Tau35 expression in mice recapitulates key features of human disease, including progressive increase in tau phosphorylation, along with cognitive and motor dysfunction. The appearance of aggregated tau suggests that Tau35 may have structural properties distinct from those of other tau species that could account for its pathological role in disease. To address this hypothesis, we performed a structural characterization of monomeric and aggregated Tau35 and compared the results to those of two longer isoforms, 2N3R and 2N4R tau. We used small angle X-ray scattering to show that Tau35, 2N3R and 2N4R tau all behave as disordered monomeric species but Tau35 exhibits higher rigidity. In the presence of the poly-anion heparin, Tau35 increases thioflavin T fluorescence significantly faster and to a greater extent than full-length tau, demonstrating a higher propensity to aggregate. We used atomic force microscopy, transmission electron microscopy and X-ray fiber diffraction to demonstrate that Tau35 aggregates are morphologically similar to previously reported tau fibrils but they are more densely packed. These data increase our understanding of the aggregation inducing properties of clinically relevant tau fragments and their potentially damaging role in the pathogenesis of human tauopathies.

## Introduction

Tauopathies are a heterogeneous group of neurodegenerative diseases that are characterized by accumulation of abnormal inclusions of the tau protein in the brain. These tau-containing aggregates form neurofibrillary tangles and other intracellular deposits, that are associated with synaptic dysfunction and neurodegeneration.^[1]^

Tau is an intrinsically unstructured protein, that binds and stabilizes microtubules. This interaction is critical for maintenance of axonal integrity in neurons.^[1]^ Six tau isoforms have been found in the human CNS which are obtained by alternative splicing of the *MAPT* gene. ^[1]^ These isoforms contain 0, 1 or 2 different 29 amino acid inserts in the protein N-terminal region, named 0N, 1N, or 2N. Each tau species contains either 3 or 4 microtubule binding repeats, generating 3R or 4R isoforms. The repeats, together with their flanking regions, are essential for microtubule binding and, in pathology, influence tangle formation.^[2-6]^ Accumulation of 3R and 4R tau isoforms results in distinct clinical manifestations: tangles comprised of paired helical filaments in Alzheimer’s disease brain contain a mixture of 3R and 4R tau, whereas 4R isoforms predominate the inclusions in progressive supranuclear palsy and corticobasal degeneration brain.^[7]^ Pick bodies are predominantly formed of 3R tau isoforms.^[8]^

The complexity of tau species found in the brain extends beyond these six isoforms. The precise nature of the tau deposits present varies among the different tauopathies, but they all contain also post-translationally modified tau species, including high phosphorylation and truncation.^[8]^ In tauopathies, phosphorylated tau forms aggregates, which are key neuropathological signatures of these disorders.^[9]^ Abnormally phosphorylated tau re-distributes from its normal localisation in axons to cell bodies and dendrites, affecting neuronal function in tauopathy brain.^[10]^ Proteolytically cleaved tau fragments are also present in human tauopathy brain, some of which may have a causal role in tau pathophysiology.^[11-15]^

We identified in human tauopathy brain a 35 kDa C-terminal tau fragment (termed Tau35), which spans residues E187 to L441 of 2N4R tau.^[15]^ Tau35 lacks the N-terminal half of tau but contains all four microtubule-binding repeats and has an intact C-terminus. We showed that even a minimal amount of Tau35 expressed in mice is sufficient to recapitulate key features of human tauopathy.^[16]^ Unlike most other tau transgenic mouse lines, the amount of Tau35 expressed in mice is less than 10% of total tau, which more accurately represents the situation in human tauopathy. Notably, despite its low expression in transgenic mice, Tau35 provokes an age-related deterioration in cognitive ability, including deficits in the Morris water maze, and an aberrant motor phenotype, which progressively worsen with age.^[16]^ Increased tau phosphorylation and pathology develop in the hippocampus and cortical regions of Tau35 mouse brain in parallel with behavioral and motor deficits. Tau35 transgenic mice also exhibit impaired short-term synaptic plasticity, possibly related to changes in membrane-associated receptors.^[17]^ These observations give rise to the hypothesis, tested here, that Tau35 has structural properties distinct from other tau species, that could make it more aggregation-prone and thereby account for its pathological behavior in disease. However, to date, no Tau35 characterization has addressed this hypothesis experimentally.

To gain new insight into Tau35 at the molecular level, we performed a detailed study of the structural and aggregation properties of Tau35. We used a combination of biophysical and biochemical techniques, including small-angle X-ray scattering (SAXS) coupled to size-exclusion chromatography (SEC), to determine the molecular weight and describe the conformational ensemble of Tau35 in solution. We also characterized the heparin-induced aggregation properties of Tau35 in comparison with the 2N3R and 2N4R tau isoforms.

We conclusively demonstrate that Tau35 is, in solution, in a stable monomeric state with a disordered random coil conformation, similar to 2N3R and 2N4R tau. The conformational ensemble of Tau35 however, has a lower degree of flexibility than intact tau. In the presence of heparin, a polyanion commonly used in the majority of *in vitro* studies of tau aggregation,^[18-21]^ Tau35 forms insoluble aggregates and has a conformational transition towards β-rich fibrillar structures that share the cross-β structure typical of paired helical filaments and all amyloid fibrils.^[22]^ Furthermore, the kinetics of self-assembly of Tau35 in the presence of heparin are faster, and Tau35 aggregates to a greater extent, than 2N4R tau. Tau35 also forms filaments that are longer and apparently more abundant than those of intact 2N4R tau. Taken together, these data provide important structural information that underpins our understanding of how tau fragmentation leads to pathogenesis in human tauopathy.

## Results

### SEC suggests preliminary evidence for the presence of a Tau35 tetramer in solution

We first aimed to characterize freshly prepared Tau35, taking the already well characterized human full-length 2N4R tau for comparison. Recombinant His-tagged tau proteins were produced in *E. coli* and purified using nickel nitrilotriacetic acid (Ni-NTA) resin. We obtained Tau35 with a yield of 15 mg/L and 2N3R and 2N4R tau with a yield of 10 mg/L culture, with greater than 90% purity for all three proteins. The apparent molecular weights of the tau species in phosphate buffered saline (PBS) were estimated from their elution volumes from a Superdex 200 10/300 column (**Figure 1A**) in comparison to globular protein standards. The retention profile for all three tau proteins differed from the values predicted from their calculated molecular weights (Tau35, 26,809 Da; 2N3R tau, 42,603 Da; 2N4R tau, 45,850 Da).^[23]^ The apparent molecular weights estimated from the calibration curve, were approximately 104 kDa for Tau35, 229 kDa for 2N3R tau, and 238 kDa for 2N4R tau (**Figure 1B**). These results are consistent with the molecular weights of a tetramer for Tau35 and pentameric species for both 2N3R and 2N4R tau. However, given that SEC elution time depends on both the shape and the net charge of the molecule, these estimates are likely to be inaccurate. For this reason, we resorted to a more powerful approach to provide information on both the state of polydispersion and on the overall shape of Tau35 when in solution.

**Figure 1.**
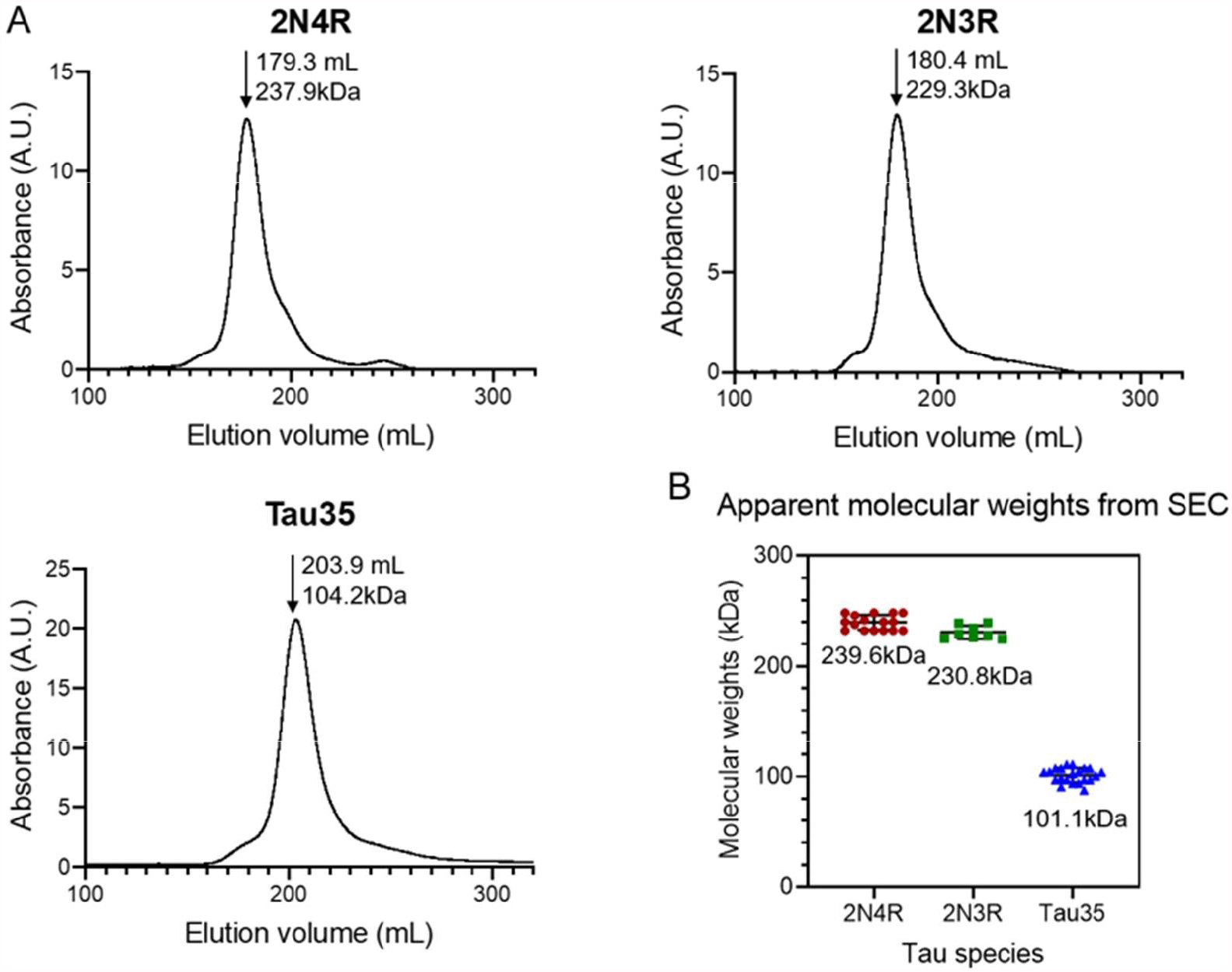
Size-exclusion chromatography of recombinant tau species. Gel filtration (Superdex 200 10/300 GL column) of recombinant 2N4R tau, 2N3R tau, and Tau35. (A) Representative elution profiles showing apparent molecular weights calculated from the elution volumes (arrows) of each tau species in comparison to a standard calibration curve. (B) Scatter plots showing the range of apparent molecular weights determined for each tau species. Bars indicate mean ± SD, n≥8.

### Characterizing the Tau35 conformation ensemble by SAXS

We used small-angle X-ray scattering coupled to size-exclusion chromatography (SEC-SAXS) to obtain accurate assessments of the molecular weights of each tau species in solution and to describe the conformational ensemble of Tau35, in comparison to that of 2N3R and 2N4R tau. SAXS is a powerful technique that can provide a low-resolution overall envelope of the molecular structure in solution.^[24]^ When coupled to SEC, SAXS enables the characterization of proteins in the absence of any contribution from potential aggregates.

The SAXS profiles and the parameters extracted from them were fully consistent with a disordered nature for all three tau species (**Figure 2A and Table 1**). This conclusion was corroborated by the monotonously increasing dimensionless Kratky plots (**Figure 2B)** and by the machine learning program DATCLASS, which classifies SAXS profiles originating from random chains.^[25]^ We obtained radii of gyration (R_g_) from Guinier analysis ^[26]^ and from the pair-distance distribution function, P(r). The Guinier approximation linearizes the SAXS intensity at low angles and relates it to the radius of gyration according to the equation: 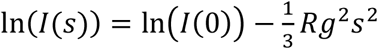. The value of Rg can then be extracted by linear fitting of a plot of ln(I(s)) vs s^2^ in the appropriate range (**Figure S1 of Supp. Info**.). An alternative estimate of the Rg can be obtained as the normalized second moment of the P(r) functions (**Figure S2 of Supp. Info**.). Lack of large deviations between these two estimates of Rg points to well-behaved SAXS curves. The Guinier plot allows also extrapolation of the forward scattering, I(0), which is proportional to the molecular weight (MW) when using concentration-normalized SAXS curves.

**Table 1.** SAXS data collection and analysis.

| **Data collection parameters** |  |  |  |
| --- | --- | --- | --- |
| Radiation source | PETRA III (DESY) |  |  |
| Beamline | EMBL P12 |  |  |
| Detector | Pilatus 6M |  |  |
| Wavelength | 0.124 nm |  |  |
| Sample-to-detector distance | 3.1 m |  |  |
| Beam size (FWHM) | 200 x 25 $\mu\text{m}^2$ | | |
| $s$ range | 0.026 – 7.36 $\text{nm}^{-1}$ | | |
| Exposure time | 1 s/frame |  |  |
| Temperature | 293.2 K |  |  |
| **Parameter (unit)** | **2N4R tau** | **2N3R tau** | **Tau35** |
| Rg<sup>a</sup> Guinier (nm) | 6.72 ± 0.28 | 6.33 ± 0.28 | 4.64 ± 0.04 |
| Rg PDDF<sup>b</sup> (nm) | 6.89 ± 0.03 | 6.32 ± 0.22 | 4.85 ± 0.02 |
| Rg from Flory scaling | 6.9 | 6.6 | 5.0 |
| Dmax<sup>c</sup> (nm) | 23.5 | 19.8 | 16.2 |
| Vp<sup>d</sup> (nm<sup>3</sup>) | 254 | 211 | 93 |
| V<sub>DAM</sub><sup>e</sup> (nm<sup>3</sup>) | 298 | 263 | 109 |
| Vc<sup>f</sup> ($s_{\text{max}}=2 \text{ nm}^{-1}$) (nm<sup>2</sup>) | 9.90 | 8.95 | 5.90 |
| MW<sup>g</sup> I(0) | 36±4 | 37±4 | 19±2 |
| MW (from Vp) | 159 | 132 | 58 |
| MW (from V<sub>DAM</sub>) (kDa) | 149 | 131 | 55 |
| MW MoW<sup>h</sup> (kDa) | 58.2 | 44.3 | 27.7 |
| MW Bayes<sup>i</sup> (kDa) | 116-151 | 93-121 | 59-66 |
| MW from sequence (kDa) | 45.8 | 42.6 | 26.8 |
| Number of amino acids | 441 | 410 | 255 |

<sup>a</sup>Rg: radius of gyration, <sup>b</sup>Rg derived from the PDDF (pair distance distribution function, see Fig.S1), <sup>c</sup>Dmax: maximal intramolecular distance, <sup>d</sup>Vp: particle volume (evaluated on the regularized intensity), <sup>e</sup>Dummy Atom Model excluded volume, <sup>f</sup>Volume of correlation, <sup>g</sup>MW molecular weight from the forward scattering I(0) (note that additional uncertainty could affect the evaluation based on the calculated E280 in the absence of any Trp residues in the sequence), <sup>h</sup>MW from Fischer's method, <sup>i</sup>MW interval with probability >90%, Bayesian method<sup>[64]</sup>. A further clarification regarding how these parameters were obtained is included in the supporting information.

**Figure 2.**
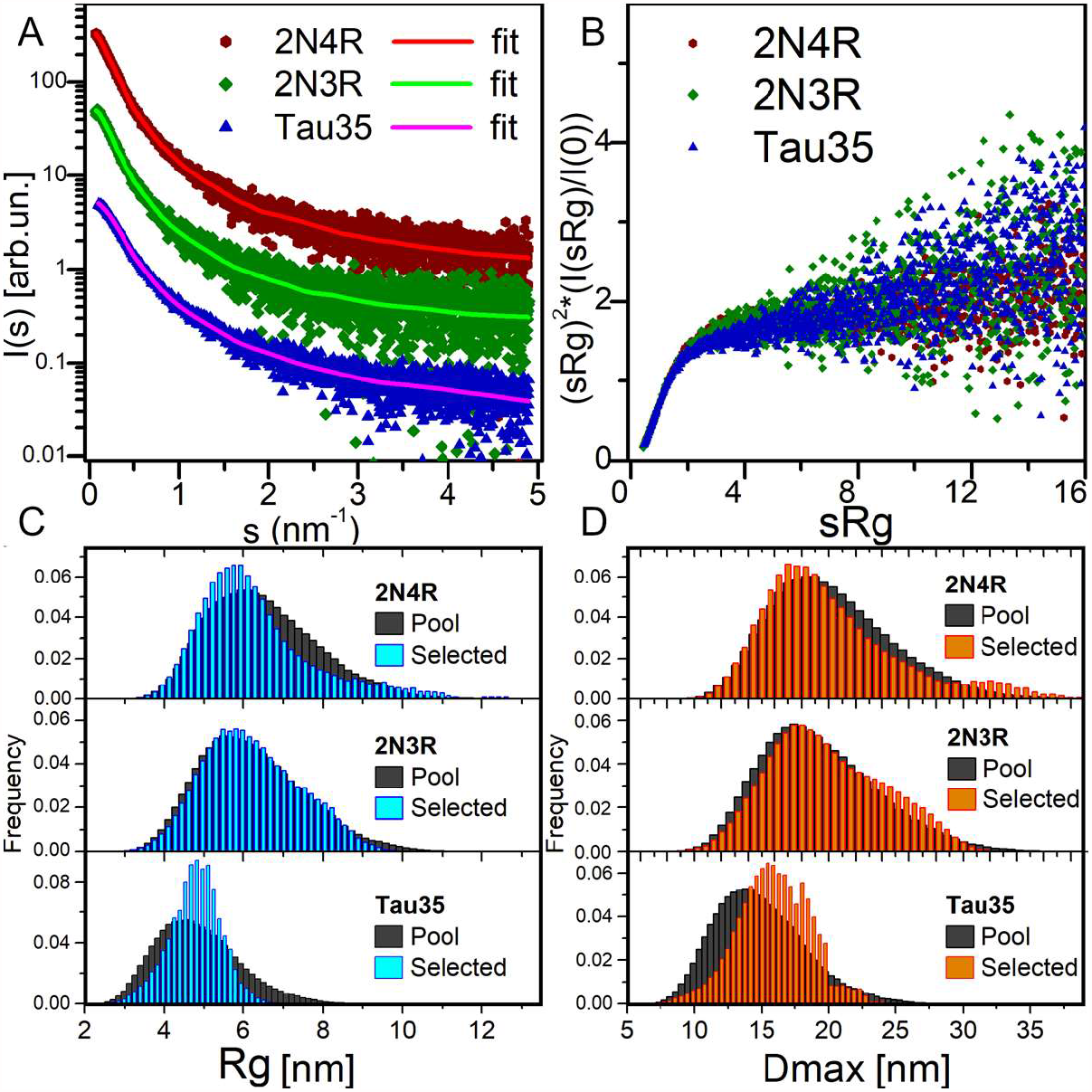
SAXS analysis. (A) SAXS profiles from SEC-SAXS for 2N4R tau, 2N3R tau and Tau35 (symbols) and corresponding typical EOM fit (solid lines, χ^2^ in range 0.98-1.00). Plots vertically displaced relative to each other for clarity. (B) Dimensionless Kratky plots for the three proteins, showing a monotonous increase, indicative of their unfolded nature. (C) Radius of gyration (Rg) distributions for the selected 2N4R tau, 2N3R tau and Tau35 ensembles compared to the random pool. (D) Maximal intramolecular distance (Dmax) distributions for the selected 2N4R tau, 2N3R tau and Tau35 ensembles.

The Rg values were in good agreement, thus providing further evidence that the SAXS curves were not affected by aggregates or higher order tau species. The Rg values were close to those predicted for chemically denatured proteins of the size of the corresponding monomeric tau species according to the Flory scaling relationship *Rg* = *R*_O_*N*^ν^ (where N is the number of amino acids, and R_0_=0.1927 nm, ν=0.588).^[27, 28]^ Applying this relationship to our constructs, we obtained theoretical Rg values of 6.9, 6.6 and 5.0 nm for 2N4R, 2N3R and Tau35, respectively. For Tau35, the Rg measured by SAXS (4.64 ± 0.04 nm from the Guinier Approximation, 4.85 ± 0.02 nm from the P(r) function) was slightly but significantly smaller than for the intact tau isoforms, pointing to a compaction likely due to increased folding or collapse in the PBS buffer used for these measurements. The R_g_ of 6.7 ± 0.3 nm for 2N4R tau was fully consistent with previous observations reporting an R_g_ value of 6.5 ± 0.3 nm.^[28]^ These results are a strong indication that Tau35, 2N3R and 2N4R tau in solution all behave as disordered monomeric species.

### Independent SAXS evidence confirms that Tau35 is a monomer

We obtained further support for these conclusions using additional independent validations. The P(r) functions of the three tau constructs provided the maximal intramolecular distance (Dmax) and an alternative estimate of R_g_, overall consistent with the value obtained by Guinier analysis. The P(r) functions were suggestive of the presence of extended tau conformations, that are typical of disordered protein ensembles (**Figure S2 of Supp. Info**.). The larger than expected sizes estimated for the tau species by comparison with similarly sized globular proteins, also differed from the molecular weights estimated from forward scattering (MW I(0)). This is an independent indication of structural disorder in the tau proteins.

Finally, we attempted *ab initio* reconstructions using the DAMMIF approach^[29]^ to provide an independent estimate of the molecular weight of the tau species in solution from the excluded volume of dummy atom models (MW_DAM_ from V_DAM_, **Table 1**). We obtained a good fit quality χ^2^ in the range of 0.9-1.2 for all models. Notably, the SAXS profiles for Tau35, 2N3R and 2N4R tau could be reasonably well fit by a simple monodisperse gaussian coil model (**Figure S3 of Supp. Info**.). ^[30]^ We also observed a lack of globular-protein-like power-law scaling in double logarithmic plots of the volume-of-correlation (Vc) as a function of amino acid sequence length (**Figure S4 of Supp. Info**.).^[31, 32]^

Taken together, these results conclusively indicate that Tau35, like 2N4R and 2N3R tau, exists as a stable monomeric species which occupies an ample conformational space.

### The Tau35 conformational ensemble is less flexible than that of 2N4R and 2N3R tau

To estimate the flexibility of the three tau ensembles, we carried out further analysis using Ensemble Optimization Method (EOM) software, which fits the data by ensembles randomly selected from pools of random conformers allowing quantitative assessment of the flexibility.^[33]^ The quality of the EOM fit to the data was excellent (χ^2^ 0.98-1.00) (**Figure 2A**, solid lines). We observed distributions similar to those of the random pools for both 2N4R and 2N3R tau (**Figure 2C, 2D**), with values of 85% and 84% for the parameter R_flex_, respectively, confirming the overall gaussian chain-like nature of these molecules. In contrast, the distribution of the selected ensemble of Tau35 was narrower than the random pool with the same number of amino acids, having a smaller R_flex_ of 73%. This indicates that Tau35 favors conformations with a narrower dispersion around a preferred size (in terms of Rg or Dmax), implying a significantly lower flexibility of this tau fragment as compared to the intact 2N3R and 2N4R human tau isoforms. It may be speculated that the relatively more ordered conformations of Tau35 could stem from the loss of a fraction of hydrophilic and charged residues, which are responsible for enhanced solvent-peptide interactions and favor disorder.

### Heparin-induced aggregation of Tau35

After having extensively characterized as-freshly-purified Tau35 and compared its behavior with full-length tau isoforms, we then turned to study the propensity of Tau35 to self-assemble by measuring its aggregation kinetics. Aggregation studies were carried out in the presence of heparin, a polyanion aggregation-inducing factor that has been extensively adopted in tau aggregation studies.^[34]^ Tau is in facts a highly soluble protein with a low propensity to aggregate spontaneously *in vitro*.^[35]^ However, early studies of 2N4R tau aggregation noted that the presence of polyanions, such as heparin or nucleic acids, can readily template tau assembly.^[18-21]^ These conditions were deemed to be closer to those observed in the cell and used in the majority of subsequent *in vitro* studies of tau aggregation.^[14, 36]^ We incubated Tau35 and 2N4R (1, 2, and 5 µM in PBS and 10 mM dithiothreitol (DTT)) at 37°C and orbital shaking at 600 rpm in the presence of heparin (at a 1:1 molar ratio). Note that the presence of a reducing agent has been demonstrated to be essential to prevent formation of disulfide bridges that would enhance filament formation.^[37]^ The kinetics of Tau35 aggregation was compared with that of 2N4R tau by monitoring the fluorescence of thioflavin T (ThT), a fluorescent probe that preferentially binds to β-rich fibrillar structures.^[38]^ Both Tau35 and 2N4R tau aggregated in a concentration-dependent manner even at low micromolar concentrations. However, Tau35 (5 µM) aggregated more than twice as fast and to a greater extent than 2N4R tau, as assessed from the slope of the aggregation curve and the higher ThT intensity at plateau, respectively (**Figure 3 and Table 2**).

**Table 2.** Heparin-induced aggregation of Tau35 and 2N4R tau in ThT assay.

| Tau concentration (μM) | Lag time (h) | | Aggregation rate (AU/h) | | Maximum intensity (AU) | |
| --- | --- | --- | --- | --- | --- | --- |
|  | 2N4R tau | Tau35 | 2N4R tau | Tau35 | 2N4R tau | Tau35 |
| 1 | 8.2 ± 0.2 | 6.8 ± 0.3 | 371 ± 21 | 411 ± 24 | 10048 ± 599 | 13569 ± 837 |
| 2 | 8.4 ± 0.2 | 6.8 ± 0.3 | 772 ± 31 | 1047 ± 53 | 17512 ± 835 | 29121 ± 1,889* |
| 5 | 7.4 ± 0.4 | 6.2 ± 0.4 | 1150 ± 62 | 2581 ± 126** | 29014 ± 1,927 | 66861 ± 3298** |

Values shown are mean ± SEM. 2N4R tau, n=8; Tau35, n=10.
\*P<0.05; \*\* P<0.01, comparison of Tau35 and 2N4R tau, Student's t-test.

**Figure 3.**
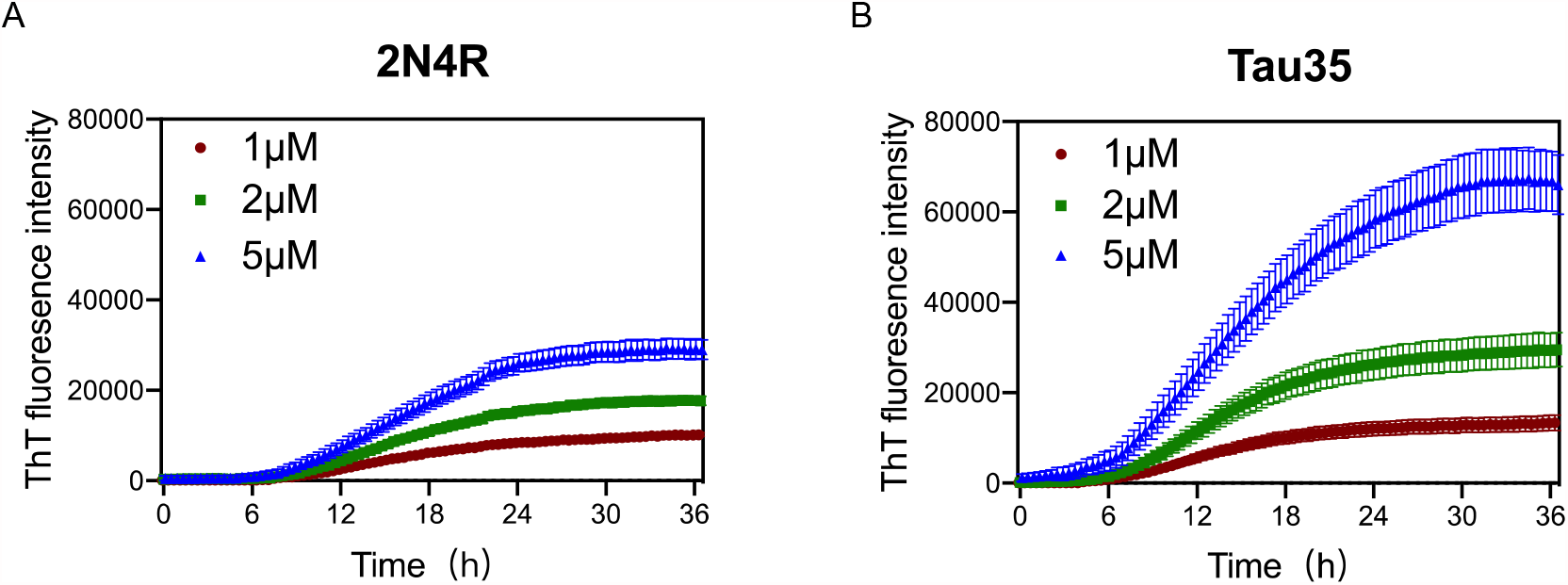
Aggregation kinetics of 2N4R tau and Tau35. Thioflavin T (ThT) fluorescence curves of 1, 2 and 5 µM (A) 2N4R tau and (B) Tau35 in PBS and DTT, following aggregation in the presence of heparin up to 36 h.

### Tau35 aggregation co-occurs with a conformational transition towards a β-rich conformation

To test whether Tau35 aggregation co-occurs with a conformational change towards a β-rich structure as observed for intact tau isoforms, we resorted to far-UV circular dichroism (CD) spectroscopy. This is a powerful method to identify the secondary structure content of a protein in solution. CD spectra of Tau35 and 2N4R tau were recorded at 20°C (**Figure 4, upper panels**). The proteins were in PBS with DTT, to which heparin was added in a 1:1 molar ratio. The CD profiles at time zero after heparin addition had features typical of a random coil structure with a single minimum at 200 nm, in agreement with established literature.^[35, 39]^ In the spectra of both Tau35 and 2N4R tau, a faint additional shoulder centered at 220 nm was also apparent and was more evident in the spectrum of Tau35. We then recorded spectra after 72 h, that is well after aggregation reaches a plateau according to the ThT fluorescence profiles. The resulting spectra remained mostly invariant in shape but the intensity of the band at 200 nm decreased likely due to precipitation or fiber deposition, with this effect being more pronounced for Tau35. To distinguish the structure of the tau aggregates from that of the soluble protein, we centrifuged the samples, separated the supernatant and re-suspended the pellet as reported previously.^[37]^ The spectra of the supernatants remained those of a random coil conformation (**Figure 4A,B**) and had features equivalent to those of the CD spectra obtained for both tau species in the absence of heparin (**Figure S5 of Supp. Info**.). In contrast, the spectra of the re-suspended pellets showed a negative band with a minimum at 230 nm, typical of a β-rich structure and indicating a conformational transition (**Figure 4C,D**). These results indicated a clear tendency of Tau35 to form β-rich fibers upon aggregation, comparable to full-length tau. Notably, following incubation of each tau protein in the presence of heparin for 72h, 69% of the starting amount of Tau35 was present in the washed pellet, whereas in comparison, only 7% of 2N4R tau was aggregated under the same conditions **(Figure 4E)**. This large increase in insoluble Tau35 after aggregation with heparin suggests that the elevated ThT fluorescence signal intensity with Tau35 is due to more efficient labeling of Tau35 fibers with ThT and/or increased relative Tau35 aggregation.

**Figure 4.**
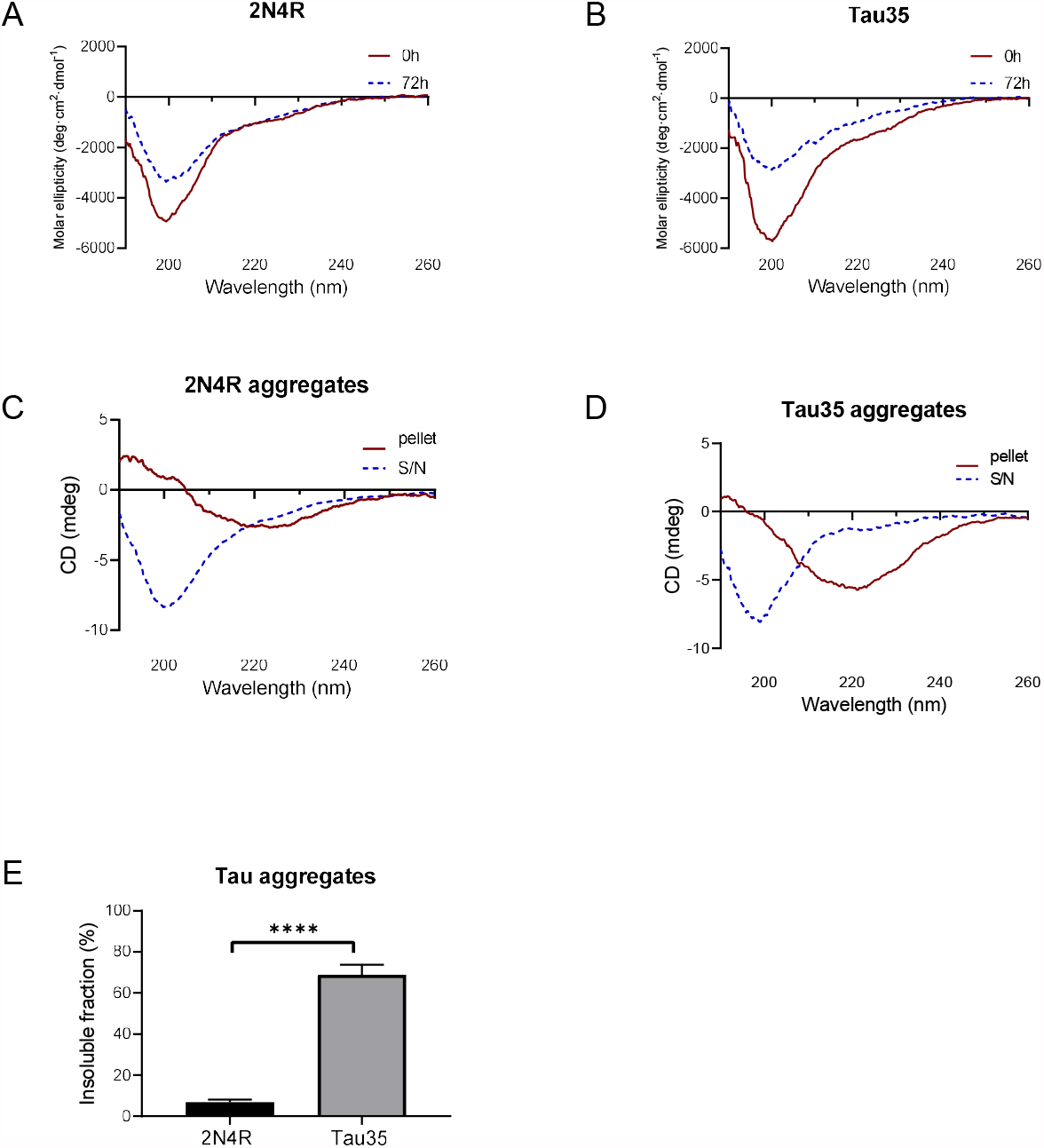
Secondary structure of 2N4R tau and Tau35 heparin-induced aggregation. Secondary structure of 2N4R tau and Tau35 heparin-induced aggregation. CD spectra obtained from (A) 2N4R tau and (B) Tau35 in 10 mM PBS at pH 7.4, following aggregation in the presence of heparin for 72 h at 37°C. (C, D) CD spectra of the same samples after centrifugation to separate out the aggregated (pellet) from the soluble (S/N) protein. (E) Graph showing the percentage of aggregated tau relative to the starting material, following heparin-induced aggregation of 2N4R tau and Tau35 for 72 h at 37°C. Bars indicate mean ± SEM, n=3, ***P<0.0001, Student’s t-test. CD spectra obtained from (A) 2N4R tau and (B) Tau35 in 10 mM PBS, pH 7.4 following aggregation in the presence of heparin for 72 h at 20°C. (C, D) CD spectra of the same samples after centrifugation to separate out the aggregated (pellet) from the soluble (S/N) protein. (E) Graph showing the percentage of aggregated tau relative to starting material, following heparin-induced aggregation of 2N4R tau and Tau35 for 72 h at 20°C. Bars indicate mean ± SEM, n=3, ***P<0.0001, Student’s t-test.

### Tau35 forms fibrillar structures that are more crowded than those formed by other tau isoforms

We compared the morphology of the heparin-induced aggregates of Tau35 and 2N4R tau by transmission electron microscopy (TEM) and atomic force microscopy (AFM). We used 100 µM Tau35 and 2N4R tau in PBS under reducing conditions following incubation with heparin for 72 h. We observed long, mostly rigid, straight filaments with some faint hint of twisting by TEM (**Figure 5A,B**). The overall features of the filaments of Tau35 and 2N4R tau were similar, but in all samples the number of aggregates was larger for Tau35, suggesting a higher tendency of this protein to form fibers under the same conditions. AFM supported similar conclusions (**Figure 5C**). The Tau35 filaments exhibited a wide range of lengths and thicknesses, ranging from approximately 80-620 nm in length and 19-28 nm in width (**Table 3**).

**Table 3.** Parameters of Tau35 fibers measured from atomic force microscopy. The values were obtained from images in which aggregate overlap would not prevent measurements for individual fibers. N was 31. SD stands for standard deviation.

| Dimension | Mean ± SD (nm) | Range (nm) |
| --- | --- | --- |
| Length | 255± 131 | 79 – 621 |
| Width | 22 ± 2 | 19 – 28 |
| Height | 5 ± 1 | 4 – 7 |

**Figure 5.**
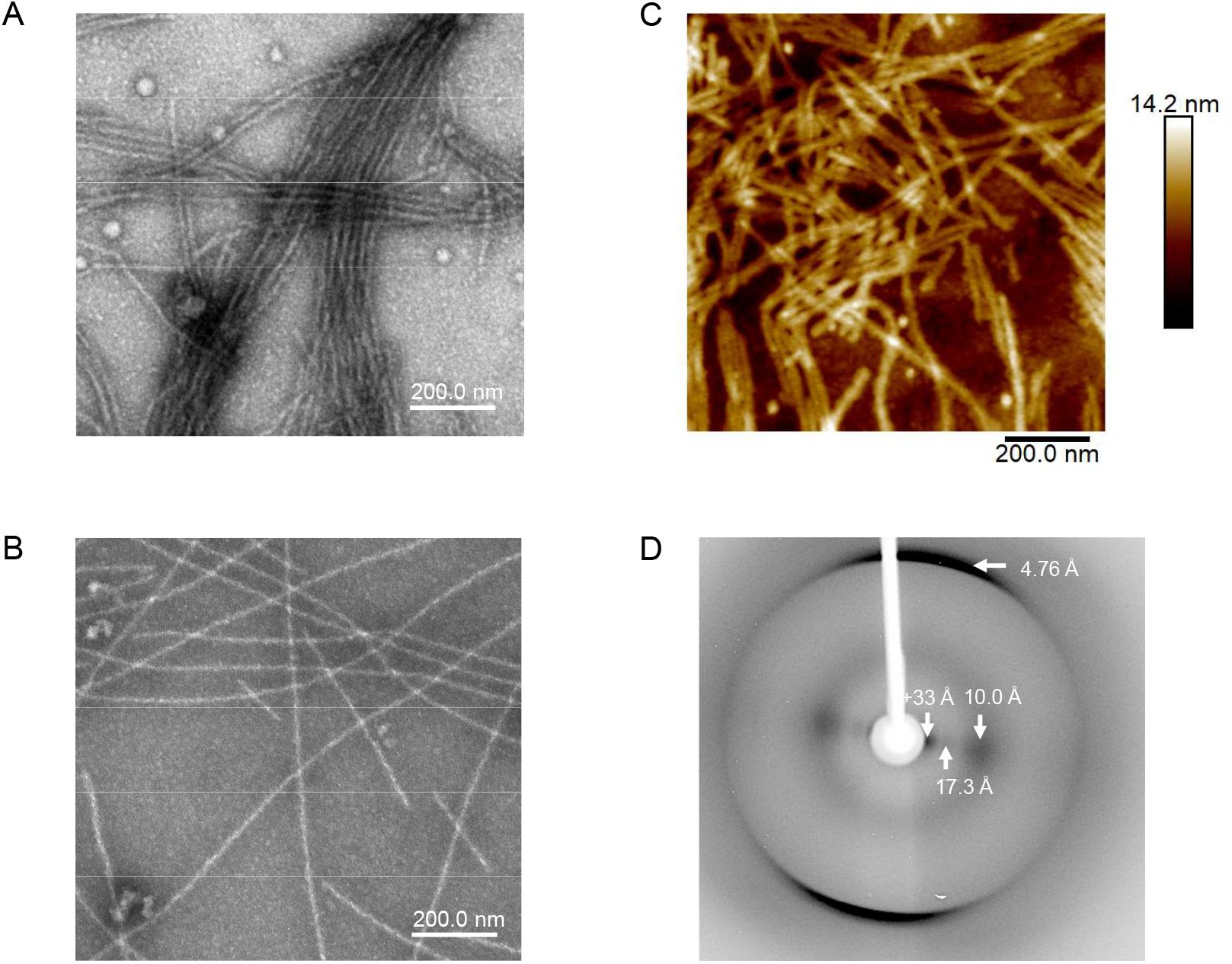
Structural and morphologic analysis of Tau35 aggregates. Electron micrographs of heparin-induced aggregates Tau35 (A) and 2N4R tau (B) incubated for 72 h at 37°C. (C) Similar morphology of Tau35 aggregates was observed by AFM. (D) X-ray fiber diffraction of aligned Tau35 fibers indicating the presence of a typical cross-β structure.

Finally, we collected data on X-ray fiber diffraction to clarify whether the Tau35 filaments have amyloid characteristics as the gold standard for detecting cross-β structures.^[40]^ A partially aligned sample of Tau35 filaments revealed the expected cross-β diffraction pattern of amyloid-like fibrils,^[41, 42]^ consistent with β-strands running perpendicular to the fiber axis with a hydrogen bonding distance of 4.76 Å and sheet spacing of about 10 Å (**Figure 5D**). Other equatorial signals at 17 and >33 Å are likely to arise from the organization of the polypeptide chain and the packing of the filaments. These data confirmed that Tau35 forms amyloid-like structures and support the role of this tau fragment as a potential driver of tau pathology in human tauopathy.

## Discussion

Tau35 is a tau fragment that has been described in human brain in progressive supranuclear palsy, corticobasal degeneration and 4R tau-predominant forms of frontotemporal dementia with Parkinsonism linked to chromosome 17, but is absent from the brains of unaffected controls, or in Alzheimer’s and Pick’s disease patients.^[15]^ Tau35-expressing mice exhibit progressive cognitive and neuropathological abnormalities including highly phosphorylated tau aggregates, that recapitulate human tauopathy.^[16]^ Tau35 has deleterious effects on cell signalling pathways that mediate pathological changes, including undergoing aberrant phosphorylation, disruption of insulin signalling and activation of the unfolded protein response.^[43]^ Despite Tau35 containing the entire microtubule-binding region of tau, the lack of the tau N-terminus results in a marked reduction in microtubule binding and defective microtubule organization in cells.^[43]^ Together, these findings suggest a mechanism through which loss of the tau N-terminus, such as in Tau35, may contribute to the pathogenesis of human disease. Based on this evidence, in the present study, we tested whether the structure and aggregation properties of Tau35 differ from those of longer, intact tau isoforms, explaining the presence of this species in tauopathies.

We used a combination of SAXS and other biophysical techniques, including fluorescence, CD, AFM, TEM and X-ray fiber diffraction to assess Tau35 and intact tau. SAXS has been used extensively to characterize 2N4R and other tau isoforms, including mutants, truncated forms and complexes,^[28, 44, 45]^ but no information is currently available for 2N3R tau or more importantly, for Tau35. Using SEC-SAXS we demonstrated that all three proteins, Tau35, 2N3R and 2N4R tau, are monomeric in solution, in agreement with previous SAXS studies on 2N4R tau.^[28]^ Since SEC-SAXS is extremely sensitive to even minute quantities of high molecular weight species, the apparently large molecular weights observed in our preliminary SEC studies are thus attributed to intrinsic disorder and elongated conformations, rather than to tau oligomerization. The soluble tau proteins have all the characteristics of intrinsically disordered monomeric proteins, with 2N3R tau having features similar to 2N4R tau, while Tau35 has more restricted motions and explores a smaller conformational space.

Further characterization of Tau35 aggregation showed that, in the presence of heparin, the protein aggregates very readily. Heparin has been customarily adopted in tau studies as an agent that conveniently enhances protein aggregation. These properties can easily be explained since addition of negatively charged heparin to positively charged tau results in the polyanion inducing coacervation and phase separation of the protein.^[46]^ Interestingly, we observed that, as compared to 2N4R tau, Tau35 has faster aggregation kinetics in the presence of heparin, possibly due the lack of the acidic, positively charged N-terminal half of tau. TEM and AFM indicated the presence of elongated fibers in aggregates of Tau35 and 2N4R tau, both which exhibited the features expected for amyloid aggregates, including cross-beta structure as proven by fiber diffraction. Tau35 fibers appear, however,more densely packed than those observed for 2N4R tau. ThT fluorescence intensity in the presence of Tau35 is also appreciably higher, suggesting increased aggregation and/or more efficient reactivity with the fluorescent probe than 2N4R tau, despite their similar structure.

In summary, the Tau35 fragment presents distinct biophysical features in comparison with intact tau, including a marked increase in its propensity to aggregate. These findings may explain some of the observations obtained in human tauopathies, in which this tau species is present in the brain, as well as in animal and cell disease models. Our results should also be put in the frame of studies of other tau fragments. For example, a truncated form of Tau35 comprising residues 297-391 (termed dGAE) and encompassing only the third and fourth microtubule binding repeats^[47]^ forms part of the proteolytically stable core of tau in neurofibrillary tangles. The dGAE tau fragment can self-assemble spontaneously *in vitro* under physiological conditions to form paired helical filament-like fibers in the absence of anionic cofactors. Moreover, assembled dGAE is more toxic than soluble, non-aggregated tau, whereas the soluble form of dGAE is internalized more readily by cells. Once inside the cell, soluble dGAE associates with endogenous tau, resulting in increased phosphorylation and aggregation of endogenous tau, which accumulates in lysosomal/endosomal compartments.^[48, 49]^ Several other N-terminally truncated tau fragments have been identified in human tauopathy brain.^[50-52]^ Notably, the N-terminal region has been reported to inhibit aggregation of full-length tau through an interaction with the C-terminal region ^[51]^, supporting our finding that Tau35, which lacks the N-terminal half, has a higher propensity to aggregate than intact tau. Furthermore, synthetic tau fragments expressed in cells show markedly differing abilities to aggregate following templated seeding by oligomeric tau from Alzheimer’s disease brain. ^[14]^ Removal of 150 residues from the N-terminus of tau resulted in enhanced tau aggregation, which was not evident following deletion of the first 50 amino acids, thereby confirming a role for the extended N-terminal region of tau in preventing its aggregation. These data suggest that tau cleavage is a critical post-translational event in the pathogenesis of tauopathy. Comparison between these results and those reported in the present work exemplifies the diverse behavior of different tau fragments that might contribute to tau pathology, which is important for understanding the structural basis of tauopathy. Our data thus add an additional layer of complexity to our understanding of tau-related species that could have implications for tau degradation and clearance in tauopathies. Further studies will be needed to determine whether this increased propensity of Tau35 to aggregate is influenced by disease-associated phosphorylation and whether this impacts on the spread of tau pathology in the tauopathies.

### Experimental procedures

Unless otherwise specified, all solvents, materials and reagents were purchased from Sigma-Aldrich.

#### Protein production

*E. coli* BL21 (DE3) pLysS strain (New England Biolabs) was used to express Tau35, *E. coli* BL21 (DE3) strain (New England Biolabs) was used for human 2N3R and 2N4R tau. The inserted plasmids contained the DNA coding the protein sequences preceded by a 6xHis tag and a tobacco etch virus (TEV) protease cleavage site at the carboxyl terminus of each protein. Protein expression was induced using 1 mM isopropyl β-D-thiogalactopyranoside at 18°C for 2 h. Bacterial pellets were re-suspended in IMAC buffer (20 mM Tris-HCl, 150 mM NaCl, 5 mM imidazole, pH 8.0) containing 10 μg/mL DNaseI and 100 μg/mL lysozyme and stored at −20°C. Thawed bacterial pellets were sonicated (4 min at output power 4, 40 cycles, Sonifier 250), heated at 100°C for 10 min and centrifuged at 60,000 g for 30 min at 4°C. All subsequent purification steps were performed at 4°C. Super nickel nitrilotriacetic acid (Ni-NTA) affinity resin (Proteinark) was mixed with the crude bacterial supernatant for 10 min before transferring to a column and washing with 10 column volumes of IMAC buffer, followed by sequential washes with 10 mM and 25 mM imidazole in IMAC buffer. His-tagged tau protein was eluted with 100 mM imidazole in IMAC buffer and incubated with TEV protease (1:10, TEV:protein) at 4°C for 16 h to remove the 6xHis tag. The protein mixture was separated by SEC and fractions containing recombinant tau were concentrated and stored at −80°C. Protein concentration was measured by Nanodrop absorption at 280 nm. Protein purity (>90%) was determined by 12% SDS-PAGE.

#### SEC measurements

Analytical SEC was carried out using a Superdex 200 10/300 GL column (Cytiva), pre-calibrated using a gel filtration marker kit (WMGF200) based on globular proteins. 10 mL recombinant Tau35, 2N3R and 2N4R tau in IMAC buffer were loaded onto the Superdex 200 column and eluted with PBS at a flow rate of 2.5 mL/min for 128 min. Protein elution profiles were monitored from the UV absorbance at 280 nm and 230 nm. Molecular weights were determined from the calibration curve of standard elution volumes.

#### SAXS measurements

Tau35, 2N3R and 2N4R tau were analyzed using SEC-SAXS ^[53]^ at the EMBL P12 beamline of the Petra III synchrotron (DESY, Hamburg, Germany).^[54]^ The protein samples were separated on a Superdex 200 Increase 10/300 GL size-exclusion column (Cytiva), monitoring the UV absorbance at 280 nm. For each sample, 100 µL protein (8-10 mg/mL) in PBS was injected onto the column and eluted with 3% (v/v) glycerol in PBS at a flow rate of 0.5 mL/min for 50 min. The eluate was flowed through a 0.9 mm borosilicate capillary, collecting 3000 SAXS frames with 0.995 s exposure time during the run. The intensity was calibrated to absolute scale using the scattering of water at 20°C. Data reduction was performed with SASFLOW.^[55]^ Buffer subtraction was conducted manually using CHROMIXS^[56]^ by selecting sample frames corresponding to the top of the main elution peak and appropriate buffer frames. The analysis yielded 1D SAXS intensity profiles I(*s*) of the species of interest, as a function of the modulus of the momentum transfer 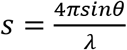 (where *2θ* is the scattering angle, λ the X-ray wavelength). SAXS data were further processed and analyzed using PRIMUS^[57]^ and the ATSAS suite.^[58]^ Concentration normalization was performed to estimate the molecular weight from the forward scattering *I(0)*, dividing the absorbance at 280 nm in correspondence of the selected sample frames by extinction coefficients calculated from the protein sequence. Pair-distance distribution functions were calculated from the SAXS profiles using the program GNOM.^[59]^ DAMMIF was used for the calculation of preliminary *ab initio* models.^[29]^

To characterize the samples as ensembles of disordered polypeptides, we fitted the SAXS profiles using EOM,^[33]^ which fits an ensemble selected from a random pool of conformations to the data, to obtain a distribution of size parameters (e.g., R_g_, Dmax), representative of the conformational space spanned by the proteins in solution. A useful parameter for the quantitative description of the flexibility is R_flex_, with a value of 100% for a fully flexible system and 0% for a rigid one. This parameter is calculated from the distributions of the size parameters obtained, treated as probability density distributions. R_flex_ is defined as the information entropy with sign inverted. Broad/uniform distributions have large R_flex_, whereas narrow distributions (corresponding to conformationally rigid systems) have R_flex_ values tending to 0. Thus, a disordered protein exploring several conformational states will have a larger R_flex_ value than a folded protein exploring a limited set of conformations in solution. For ensemble analysis, the program EOM 2.0 was used.^[60]^ For each construct, 10,000 random structures were generated based on the known (monomeric) amino acid sequence and five repeating EOM selection runs were carried out. No intra-molecular restraints were imposed. Fitting to a monodisperse gaussian coil model was performed by the program SasView 5.0 (SasView Version 4.2, Zenodo). The SAXS data and typical EOM fits for Tau35, 2N3R and 2N4R were submitted ot the SASBDB with entry codes SASDLS4, SASDLT4 and SASDLU4 respectively.

#### ThT fluorescence assays

Aggregation kinetics for all constructs were determined using ThT-binding assays in Greiner Bio-One CELLSTAR plates and monitored using a BMG FLUOstar Omega microplate reader. Proteins were freshly prepared and filtered (0.2 µm) before each assay to ensure removal of any contingent pre-formed oligomers. Samples were diluted in PBS and 10 mM DTT to final concentrations of 1, 2, and 5 µM protein, in the presence of 1:1 molar ratios of heparin (porcine intestinal mucosa, H4784, Sigma-Aldrich), followed by addition of ThT to a final concentration of 20 µM. Protein aggregation was induced at 37°C with orbital shaking at 600 rpm and the fluorescence intensity of ThT was recorded (excitation/emission, 430/485 nm) for up to 72 h. The reported data derive from at least eight repetitions of each experiment.

#### In vitro tau aggregation for CD spectroscopy and fiber morphology assays

Heparin was added in a 1:1 molar ratio to 100 µM Tau35 or 2N4R tau in PBS containing 10 mM DTT. Protein aggregation was induced at 37°C with orbital shaking at 600 rpm for 72 h. For some experiments, tau was aggregated in pre-weighed tubes and following aggregation, samples were centrifuged at 20,000 g for 30 min at 4°C to separate aggregated from soluble tau. The pellets containing insoluble tau were washed with ultrapure water and centrifuged as above. The washed pellets were allowed to air-dry overnight at ambient temperature before measuring their mass to determine the molar amount of tau present in each pellet. Aggregated tau was subjected to CD spectroscopy. Fiber morphology was assessed by AFM, TEM and X-ray diffraction, as described below.

#### CD spectroscopy

Far-UV CD spectra of soluble tau species were recorded at 20°C using a 0.1 mm path-length quartz cuvette under a constant nitrogen flush at 2.0 L/min (JASCO-1100 or J715 spectropolarimeters). Spectra of heparin-free Tau35 and 2N4R tau were acquired as the sum of 10 scans recorded. The spectra obtained from heparin-treated samples as described above, were collected after centrifuging the sample at 20,000 g for 30 min at 4°C to separate the fibrils from the soluble fraction. The pellet was washed and re-suspended in 10-50 μL 0.22 μm filtered Milli-Q water. CD spectra of the pellets and supernatants were measured at 20°C. The spectra obtained from the soluble proteins were corrected for the buffer signal and known concentration, expressing as mean residue molar ellipticity, θ (deg*cm^2^*dmol^−1^). All spectra were collected in triplicate.

#### Fiber morphology by AFM

Height and peak force error AFM images were obtained on a Bruker Multimode 8 microscope with a NanoScope V Controller (Bruker UK Ltd). Images were acquired operating in peak force tapping mode using ScanAsyst Air cantilevers using ScanAsyst probes with a 2 nm nominal tip radius of curvature. Image data were obtained at a peak force frequency of 4 kHz and a line rate of 3 Hz, at a resolution of 512 pixels/line. 10 µL sample was loaded onto freshly cleaved mica and incubated for 10 min at ambient temperature. The liquid excess was dried off from the mica, which was rinsed three times with a gentle flux of 0.22 μm filtered Milli-Q water. Aggregated Tau35 was left on formvar/carbon-coated 400-mesh copper grids (Agar Scientific) for 60 s before blotting the excess using a filter paper. The grids were washed with 4 μL 0.22 μm filtered Milli-Q water and negatively stained with 4 μL 2% (w/v) uranyl acetate for 30 s. TEM images were collected using a JEOL JEM1400-Plus transmission electron microscope operated at 80 kV. Detection was achieved using a 4kx4K OneView camera (Gatan). Acquisitions were performed at 25 fps and automatically corrected for drift using DigitalMicrograph® software (GMS3, Gatan).

#### X-ray fiber diffraction

Aggregated tau samples were centrifuged at 13,000 g for 10 min at ambient temperature, washed three times with 0.22 μm filtered Milli-Q water and re-suspended in 10 mL water. A droplet of approximately 10 μL was suspended between two capillary tubes with wax tips and allowed to dry at ambient temperature sealed within a Petri dish, as previously described.^[61]^ The sample was mounted on a goniometer head and diffraction data were collected using a CuKα Rigaku rotating anode and Saturn CCD detector with exposure times of 30-60 s and specimen to detector distances of 50 and 100 mm. Diffraction patterns were inspected using iMosflm ^[62]^ and converted to tif format. Diffraction spacings were measured using CLEARER software.^[63]^

## Abbreviations

AFM: atomic force microscopy
CD: circular dichroism
DTT: dithiothreitol
EOM: Ensemble Optimization Method
Ni-NTA: nickel nitrilotriacetic acid
PBS: phosphate buffered saline
SEC: size-exclusion chromatography
TEM: transmission electron microscopy
TEV: tobacco etch virus
ThT: thioflavin T

## Acknowledgements

We wish to thank Dr Alessandro Sicorello for technical assistance during the early stages of this project.

## Funding and additional information

The work was funded by the Dementia Research Institute (RE1 3556). The work benefited from the use of the SasView application (http://www.sasview.org/), originally developed under NSF Award DMR-0520547. SasView also contains code developed with funding from the EU Horizon 2020 programme under the SINE2020 project Grant No 654000. CL is funded by a King’s College London-China Scholarship Council studentship. The authors acknowledge the electron microscopy imaging centre of the University of Sussex, funded by the School of Life Sciences, the Wellcome Trust and the RM Phillips fund, for their support and assistance with this study. D.Sv. and S.D. acknowledge funding from the Joachim Herz Stiftung (Hamburg, Germany), grant “Biomedical Physics of Infection”.

## Conflict of interest

The authors declare that they have no conflicts of interest with the contents of this article.

